# Regulation of multidrug efflux pumps by TetR family transcriptional repressor negatively affects secondary metabolism in *Streptomyces coelicolor* A3(2)

**DOI:** 10.1101/2022.11.11.516236

**Authors:** Yukun Lei, Shumpei Asamizu, Takumi Ishizuka, Hiroyasu Onaka

## Abstract

*Streptomyces* spp. are well-known producers of bioactive secondary metabolites (SMs) that serve as pharmaceutical agents. In addition to their ability to produce SMs, *Streptomyces* spp. have evolved diverse membrane transport systems to protect cells against antibiotics produced by itself or other microorganisms. We previously screened mutants of *Streptomyces coelicolor* that show a phenotype of reduced undecylprodigiosin (RED) production in a combined-culture with *Tsukamurella pulmonis*. Here, we identified a point mutation, which reduced RED production, by performing genome resequencing and genetic complementation. We found that inactivation of the *sco1718* gene encoding the TetR family transcriptional regulator (TFR) produced a deficient phenotype for several SMs in *Streptomyces coelicolor* A3(2). Electrophoretic mobility shift assay and quantitative reverse transcription-PCR experiments demonstrated that SCO1718 repressed the expression of adjacent two-component ATP-binding cassette (ABC) transporter genes (*sco1719-20*) by binding to the operator sequence in the 5′-UTR. Notably, the Δ*sco1718* mutant showed increased resistance to several antibiotics of other actinomycete origin. In the genome of *S. coelicolor* A3(2), two other sets of TFR and two-component ABC transporter genes (*sco4358-4360* and *sco5384-5382*) were found, which had similar effects on the phenotype for both secondary metabolism and antibiotic resistance. Our results imply the switching of cell metabolism to direct offence (antibiotic production) or defense (efflux pump activation) using costly and limited quantities of cell energy sources (e.g., ATP) in the soil ecosystem.

**IMPORTANCE:** The bacterial metabolic potential to synthesize diverse secondary metabolites (SMs) in the environment has been revealed by recent (meta-)genomics of both unculturable and culturable bacteria. These studies imply that bacteria are continuously exposed to harmful chemical compounds in the environment. *Streptomyces* spp. contain antibiotic efflux pumps and SM biosynthetic gene clusters. However, the mechanism by which soil bacteria, including *Streptomyces*, survive against toxic compounds in the environment remains unclear. Here, we identified three sets of TFR-ABC transporter genes in *Streptomyces coelicolor* A3(2). We found that each TFR controlled the expression of a respective ABC transporter, and the expression of all ABC transporters negatively impacted SM production and increased antibiotic resistance. Notably, bioinformatic analysis indicated that these TFR-ABC transporter gene sets are highly conserved and widely distributed in the genome of *Streptomyces* species, indicating the importance of systematic regulation that directs antibiotic production and xenobiotic excretion.

## INTRODUCTION

*Streptomyces* species are ubiquitous soil bacteria that are widely recognized as producers of antimicrobial secondary metabolites (SMs) (1, 2). A considerable number of *Streptomyces* genome sequences have revealed the presence of 20–50 diverse biosynthetic gene clusters (BGCs) for SMs in each *Streptomyces* genome, which indicates a major untapped potential to elucidate SM biosynthesis (3, 4). Several strategies have been developed to unveil the metabolites encoded by the cryptic BGCs (5, 6); however, only a limited number of BGCs have been expressed under laboratory culture conditions, and the most regions remain unknown. A large number of diverse and several conserved BGCs in these soil bacteria may be advantageous for survival in the natural environment, including competition for nutritional resources with other microorganisms (7).

SM production by *Streptomyces* involves complex cell metabolism, and some processes are strictly linked and regulated by cell differentiation (8) as well as induced by environmental stresses such as cell envelope damage, osmotic stress, and oxidative stresses (6, 7). Although several dedicated (clustered) systems involved in SM production have been characterized in detail, such as SARP (9), random mutagenesis experiments in studies have frequently revealed “orphan” (standalone) genes, which affect the production of SM, presumably in an indirect manner (10, 11). Determining the role of these orphan genes in the production of SMs is difficult; however, elucidating the uncharacterized orphan systems that affect SM production is important for a deeper understanding of how SM production systems participate in the whole cell system.

Bacteria have evolved defense (resistance) systems to protect cells from the harmful bioactive compounds produced by other microorganisms (12). One well-known mechanism of antibiotic resistance involves efflux pumps that reduce the intracellular concentration of toxic molecules (13). These efflux pumps include the major facilitator superfamily (MFS) (14), resistance-nodulation-division (RND) family (15), small multidrug resistance family (16), multidrug and toxin extrusion family (17), and ATP-binding cassette (ABC) family transporters (18). Generally, ABC transporters consist of two to four protein components, including ATP-binding proteins, one or two membrane proteins (permease), and a substrate-binding protein (SBP) (19). ABC transporters containing SBPs function as importers; however, ABC transporters without SBP can function as importers or exporters (18, 19).

The model actinomycete *Streptomyces coelicolor* A3(2) is estimated to possess 795 putative transporter genes, of which 432 are predicted to be ABC transporter genes according to TransportDB 2.0 (20). ABC transporters are importers of diverse substrates, such as sugar (21), phosphate (22), *N*-acetylglucosamine (23, 24), and siderophores (25). ABC transporters also function as drug efflux pumps and can be classified into two types: dedicated or promiscuous when exporting substrates. ABC transporter genes located within an SM-BGC are predicted to possess substrate specificity and export the encoded SM for self-resistance, such as *cchCDEFGI* genes (*sco0491* and *sco0493-97*) in coelichelin biosynthesis (26), *sco3223-24* in calcium-dependent antibiotic (CDA) biosynthesis (27), *ramAB* (*sco6757-58*) in SapB biosynthesis (28, 29), and *sco7689-90* in coelibactin biosynthesis (30). In contrast, part of standalone ABC exporters are known to exhibit substrate tolerance and function as multidrug efflux pumps (31, 32). *S. coelicolor* A3(2) possesses hundreds of putative transcriptional regulators, and the TetR family transcriptional regulator (TFR) forms one of the largest groups comprising approximately 150 members (33). TFRs regulate a wide range of physiological processes, including carbon metabolism, quorum sensing, antibiotic resistance, cell division, and SM biosynthesis (33, 34). Several standalone ABC transporter genes exist adjacent to transcriptional regulator genes (35). Generally, TFRs function as dimers and bind to the operator sequence (palindrome sequence) to block the binding of RNA polymerase to the promoter (34). When specific small-molecule ligands bind to the TFR, conformational changes are induced in TFR and the ability to bind to the DNA is lost, which allows the binding of RNA polymerase and transcription of downstream genes (34).

Co-culture involving actinomycetes and mycolic acid-containing bacteria (e.g. *Tsukamurella pulmonis* TP-B0596) termed “combined culture” has been performed to induce SM production in actinomycetes (36-38). To date, 40 new SMs have been isolated from 14 actinomycetes and this co-culture method represents one of the most successful methods for natural product discovery (compounds listed in Table S1) (39, 40). Previously, we reported the generation of an *S. coelicolor* JCM4020 mutant library using a carbon ion (^12^C^5+^) beam to screen mutants with different undecylprodigiosin (RED) production levels to identify the genes responsible for RED production during interaction with *T. pulmonis* (11). Here, we further investigated the obtained mutants and identified the TFR gene *sco1718*, in which disruption was responsible for the phenotype of reduced RED production. We report a functional analysis of a conserved system in which TFRs regulate the expression of ABC transporter multidrug efflux pumps.

## RESULTS

### Genome re-sequence of Mt-209003 and identification of point mutations

Previously, we generated a mutant library using *S. coelicolor* JCM4020 using carbon-ion beam irradiation to screen RED production-deficient mutants (11). To identify the genes responsible for RED production, we further analyzed point mutations induced in the Mt-209003 genome of strains showing a phenotype of reduced RED production by screening with *T. pulmonis* TP-B0596 interaction (Fig. S1). We re-sequenced the genome of Mt-209003 and determined the point mutations (Table S2). Among the point mutations, the JCM4020_18610 gene contained a point mutation (34delT) that caused a frame shift (Trp12fs). The product of JCM4020_18610 showed 100% identity with the *sco1718* gene product in *S. coelicolor* A3(2) (hereafter JCM4020_18610 = *sco1718*^JCM4020^). We performed genetic complementation of *sco1718*^JCM4020^ in trans under the control of the original promoter in Mt-209003, and demonstrated that RED production under combined-culture with *T. pulmonis* was rescued (Fig. S1). Thus, *sco1718*^JCM4020^ is responsible for the reduced production of RED in strain JCM4020. Therefore, we further analyzed the involvement of SCO1718 in the reduced RED production phenotype.

### Δsco1718, a TetR-like repressor-inactivated mutant, showed reduced production of RED

We used the most studied actinomycete, *S. coelicolor* A3(2) M145, for further analyses, which maintain an identical system. To confirm that inactivation of *sco1718* confers an identical phenotype in *S. coelicolor* A3(2), we generated a targeted gene deletion mutant Δ*sco1718*, which successfully showed a phenotype comparable to that of JCM4020 (Fig. 1A). Adjacent to the TFR-encoding *sco1718* gene, *sco1719-20* which encoded a two-component ABC transporter was present in the opposite direction with a 63 bp intergenic region (Fig. 2A). SCO1719 contained an ATP-binding domain and SCO1720 contained a transmembrane domain. SCO1719/20 showed similarity to DrrA/B, a dedicated doxorubicin efflux pump in *Streptomyces peucetius* (47.7/32.6% identity, Table S3), and showed similarity to a multidrug resistance ABC transporter (also called DrrA/B) derived from *Mycobacterium tuberculosis* H37Rv (41.1/24.0% identity, Table S3). Compton et al. showed that overexpression of *sco1719-20* in *S. coelicolor* A3(2) increases resistance to antibacterial acyldepsipeptide (ADEP), which indicates that SCO1719/20 functions as a multidrug resistance-conferring ABC transporter (31).

**FIG 1.**
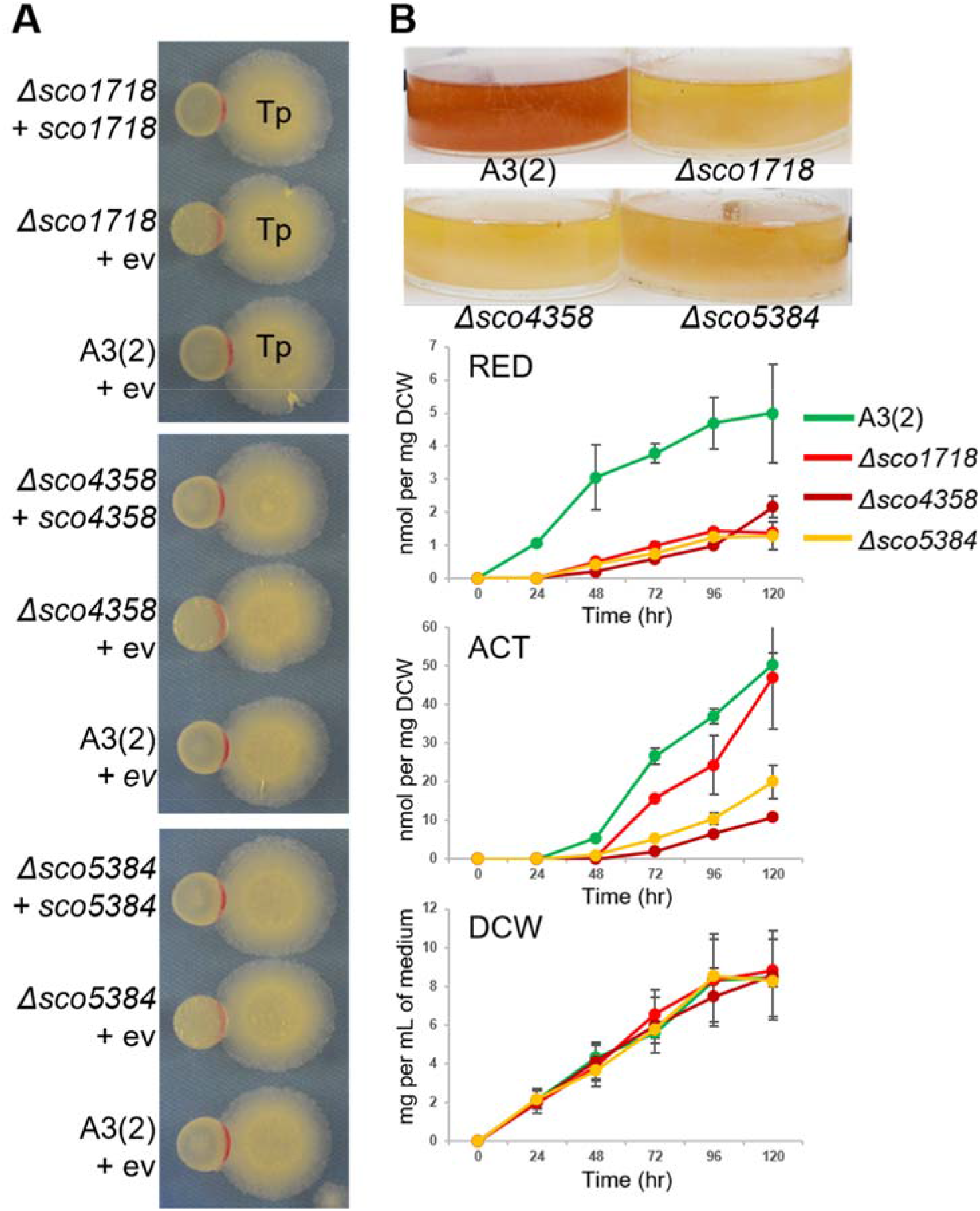
Reduced production of RED and ACT by TFR-knockout mutants. (A) Combined culture of TFR-knockout mutants (Δ*sco1718*, Δ*sco4358*, and Δ*sco5384*) with *T. pulmonis* TP-B0596. *S. coelicolor* A3(2) strains and *T. pulmonis* cell stock were spotted on YGGS agar and incubated at 30°C for 3 d. TFR-knockout strains showed reduced RED production, and genetic complementation of the respective gene rescued the RED production. ev: empty vector (pTYM1a). (B) Quantification of produced RED and ACT and the growth curve of TFR-knockout mutants in R2YE medium. Photos were taken after 48 h of culture. RED (nmol /mg DCW). ACT production (nmol / DCW). Growth curves were generated based on DCW (mg/ml of medium). Error bars correspond to the standard errors of the means for three replicates in each culture. TFR, TetR family transcriptional regulator; RED, undecylprodigiosin; ACT, actinorhodin; DCW, dry cell weight.

**FIG 2.**
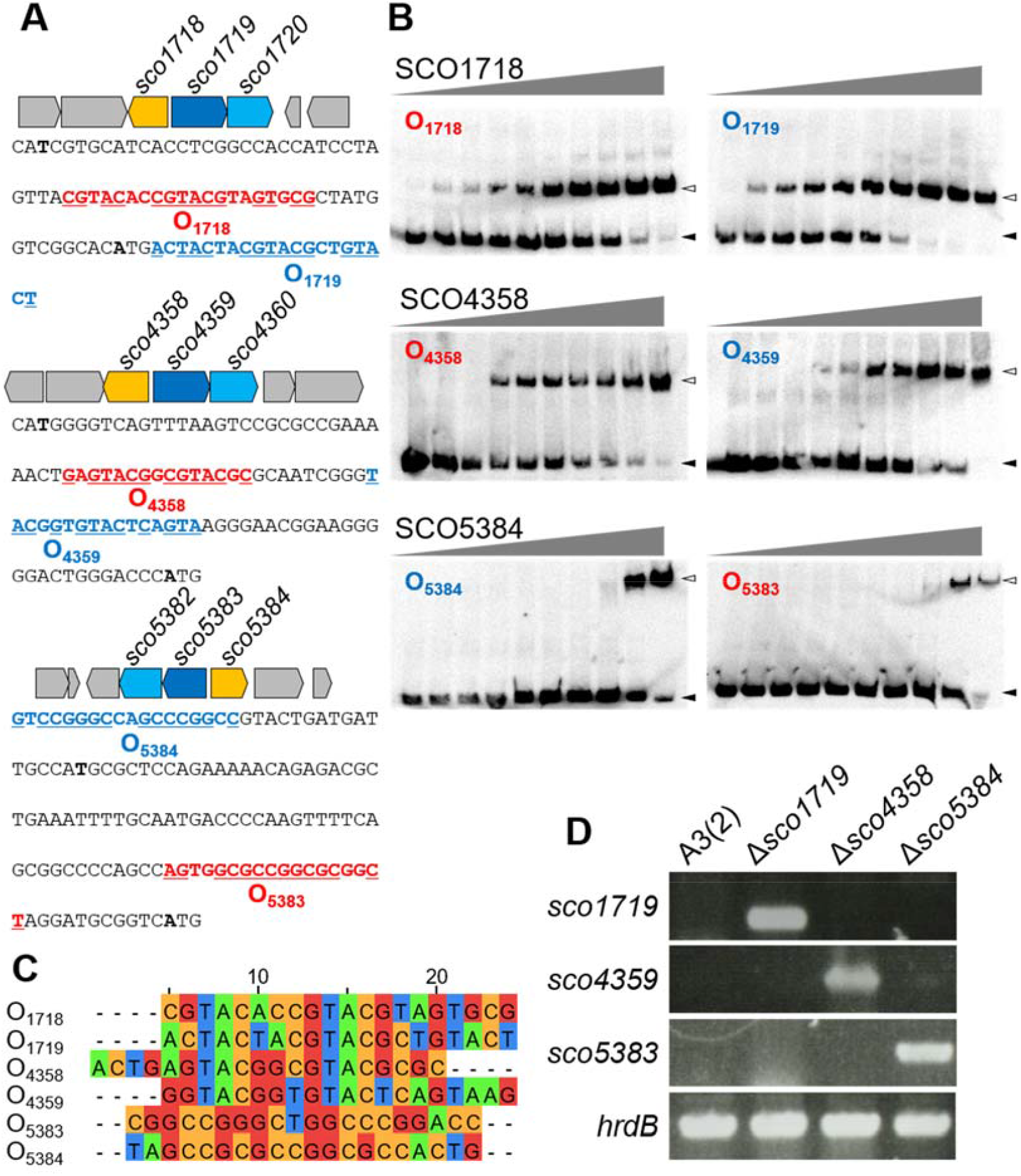
TFRs repress the expression of ABC transporters. (A) Genetic map of three TFR-ABC transporter regions in *Streptomyces coelicolor* A3(2). (B) Electrophoretic mobility shift assay with DIG-labelled oligo-DNA probes. The free probes are indicated by filled arrows and DNA–protein complexes are indicated by blank arrows. DNA probe: 0.2 nM; SCO1718: 0, 0.2, 0.5, 1, 2, 5, 10, 20, 50 nM; SCO4358 and SCO5384: 0, 0.5, 1, 2, 5, 10, 20, 50 100 nM. (C) Sequence alignment of operator regions. (D) Semi-quantitative PCR of ABC transporter genes in TFR-inactivated mutants. *hrdB* gene was used as an internal control. PCR amplification: 25 cycles. TFR, TetR family transcriptional regulator.

*S. coelicolor* A3(2) possesses approximately 432 ABC transporter genes, which have been predicted via TransportDB 2.0 (20). Using DrrA (Rv2936) from *M. tuberculosis* as a query, we searched for homologs using Kyoto Encyclopedia of Genes and Genomes (KEGG) database. In total, 31 putative proteins showed >100 bit scores (Table S4), and flanking regions were analyzed using Clinker (41) and aligned according to phylogenetic analysis using MEGA-X (42) (Fig. S2). To further elucidate the function of the unique gene set consisting of TFR and two-component ABC transporter genes comprising ATP-binding protein gene and transmembrane protein gene in adjacent, we searched for other gene sets similar to *sco1718-20*. Two other sets, *sco4358-60* and *sco5382-84*, were successfully identified (Fig. 2A and Fig. S2). The other two gene sets were also standalone and did not cluster with the putative SM-BGC.

### TFRs function as repressors of ABC transporter

TFR generally forms dimers and is known for its autoregulation (34). Upon searching for the operator sequence in all three gene sets, two palindrome sequences were predicted in each of the 5′-untranslated region (5′-UTR) of the TFR and ABC transporter genes (Fig. 2A). An electrophoretic mobility shift assay (EMSA) was performed to identify whether the proteins recognized the respective palindromic sequences as binding sites. The TFR genes (*sco1718, sco4358*, and *sco5384*) were cloned and overexpressed under control of the T7 promoter in *E. coli* BL21(DE3) and purified as *N*-terminal His-tagged proteins using Ni-NTA resin (Fig. S3). The recombinant proteins were mixed with the respective oligo-DNA probes containing palindromic sequences from the 5′-UTR of the three TFR genes (Table S5). By adding each TFR protein, a specific shift band of the DNA probes was successfully observed in the EMSA assay (Fig. 2B). We further introduced base substitution mutations to disrupt the recognition sequences. Incubation with the mutations did not result in a shifted band in the EMSA assay, confirming the importance of the specific sequence for recognition by respective TFRs (Fig. S4). Next, to confirm that the three TFRs (SCO1718, SCO4358, and SCO5384) repress the expression of each adjacent ABC transporter, we analyzed transcription levels via semi-quantitative PCR. In addition to Δ*sco1718*, knockout mutants of *sco4358* and *sco5384* were generated via genome editing using the clustered interspaced short palindromic repeats (CRISPR)-cas9 system. The parent strain A3(2) and the three mutants were cultured in R2YE medium for 24 h, and total RNA was extracted. cDNA was generated from isolated total RNA, and specific primers (Table S6) were used to amplify the ABC transporter gene regions of *sco1719, sco4359*, and *sco5383*. The results clearly showed that the transcription of each ABC transporter gene was significantly increased in the respective TFR-knockout mutants (Fig. 2D). Notably, although the operator sequences of *sco1718* (O_1718_) and *sco4358* (O_4358_) possessed sequence similarity (Fig. 2C), regulation of cross-talk between SCO1718 and SCO4358 was not found. For example, in the Δ*sco1718* mutant, amplification of only the *sco1719* region was observed, whereas no amplification of the *sco4359* and *sco5383* regions was observed (Fig. 2D). Similar results were observed for the Δ*sco4358* and Δ*sco5384* mutants (Fig. 2D). Therefore, the TFRs are dedicated to ABC transporters genetically adjacent to them.

### Production of SMs was decreased upon expression of ABC transporters

As the Δ*sco1718* mutant showed a phenotype of reduced RED production (Fig. 1), we further investigated the production of other SMs by TFR mutants. *S. coelicolor* A3(2) produces RED, blue-pigmented actinorhodin (ACT), CDA (27), yellow-pigmented coelimycin (cpk) (43), and desferrioxamines (DESs) as a siderophore (44). Notably, all three TFR-knockout mutants (*sco1718, sco4358*, and *sco5384*) showed reduced RED and ACT production compared to that in the parent A3(2) strain, although no significant defects in their growth were observed in the monoculture using R2YE medium (Fig. 1B). The reduced RED production phenotype was also observed even when the mutants were combined-cultured with *T. pulmonis* (Fig. 1A). This indicates that the basal level of RED (or ACT) production is decreased by the overexpression of ABC transporters. As the production of two representative SMs (RED and ACT) was decreased in all three TFR mutants, we further tested other SMs. Notably, for the production of CDA, the Δ*sco4358* mutant alone showed significantly decreased production, whereas the other two mutants did not show significant differences in production of CDA compared to that in the parent strain (Fig. 3A). For the production of cpk, similar to the results of CDA, the Δ*sco4358* mutant alone showed a decrease in production; the other two mutants did not show significant differences in the production of cpk compared to that in the parent strain (Fig. 3B). For the production of DES, the Δ*sco1718* and Δ*sco5384* mutants showed slightly reduced production of DES; however, the Δ*sco4358* mutant did not show a significant difference in the production of DES compared to that in the parent strain (Fig. 3C).

**FIG 3.**
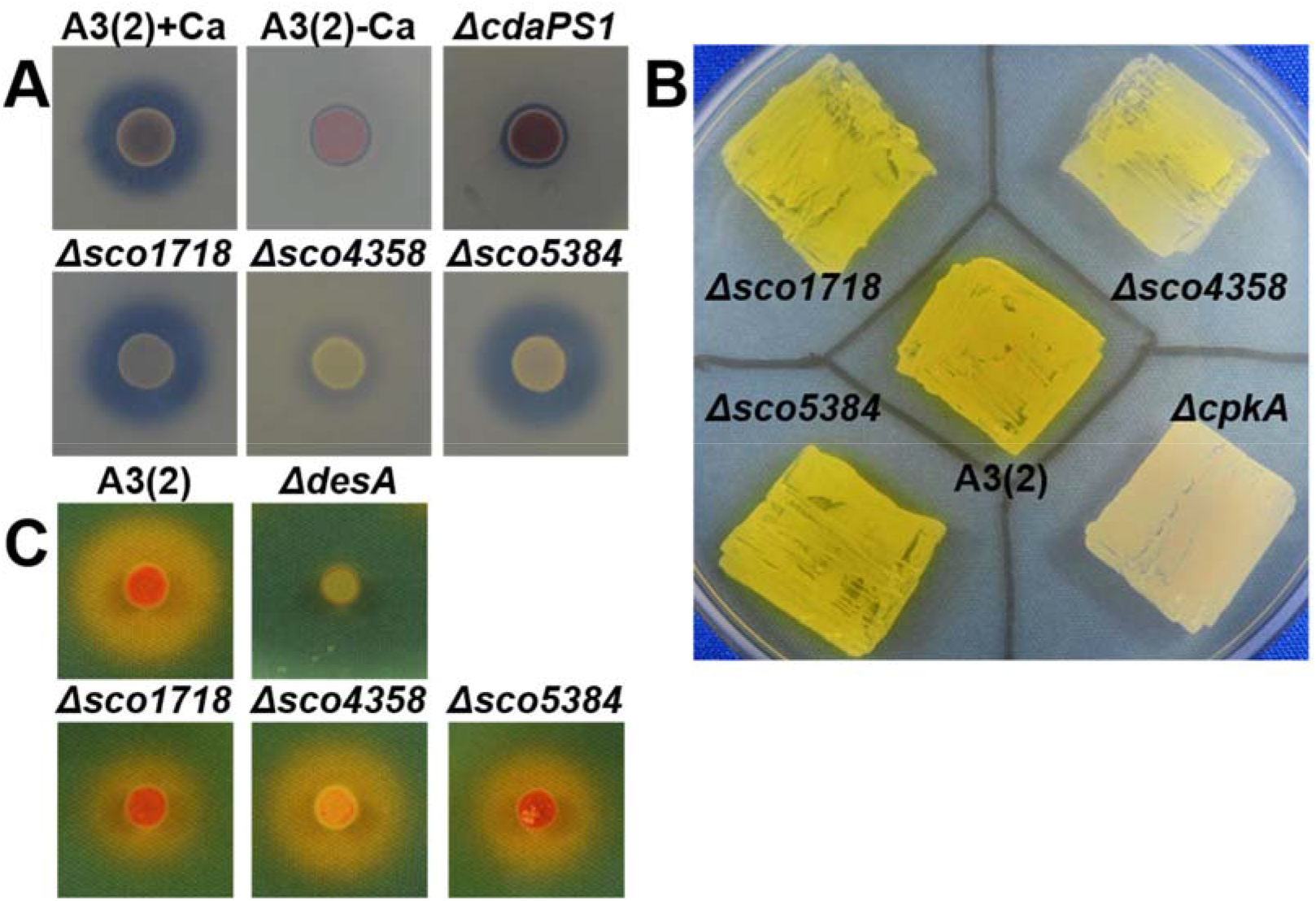
Production of secondary metabolites by *Streptomyces coelicolor* A3(2) and TFR-knockout mutants. (A) CDA production bioassay against *Bacillus subtilis* on NAHU solid medium. A plate without calcium addition and a plate spotted with non-CDA-producing mutant (Δ*cdaPS1*: CDA peptide synthase I disruptant) were used as controls. (B) Assay of coelimycin production on 79NG solid medium. Plate spotted with non-coelimycin-producing mutant (Δ*cpkA*: peptide synthase disruptant) was used as a control. (C) Assay of desferrioxamine production on R2YE solid medium. Plate spotted with non-desferrioxamine-producing mutant (Δ*desA*: desferrioxamine synthase disruptant) was used as a control. TFR, TetR family transcriptional regulator; CDA, calcium-dependent antibiotic.

### TFR-regulated ABC transporters function as multidrug efflux pumps

Gominet et al. reported that disruption of the *sclR* gene (ortholog of *sco4358*) encoding TFR in *Streptomyces lividans* 1326 confers resistance to antibacterial ADEP, and disruption of the *sclA* gene (ortholog of *sco4359*) encoding the ABC transporter confers sensitivity to ADEP(32). Subsequently, Compton et al. reported that disruption of *sco1718* and overexpression of the genetic locus *sco1719-20* in *S. coelicolor* A3(2) increases resistance to ADEP (31). Therefore, we tested the sensitivity to other antibiotics upon expression of the three standalone ABC transporters. First, the susceptibility to 19 natural product-origin antibiotics was tested using a paper disc diffusion assay for the TFR disruptants and parent strain (Table S7). Δ*sco1718* and Δ*sco5384* showed increased resistance to erythromycin, rifamycin SV, thiostrepton, goadsporin(45), and fusidic acid compared to that of the A3(2) parent strain (Fig. 4). Δ*sco4358* showed increased resistance to thiostrepton, goadsporin, and daunorubicin compared to that of the A3(2) parent strain (Fig. 4). To further confirm that resistance was a result of the expression of ABC transporters, we generated double-knockout mutants of TFR and ABC transporter genes.

**FIG 4.**
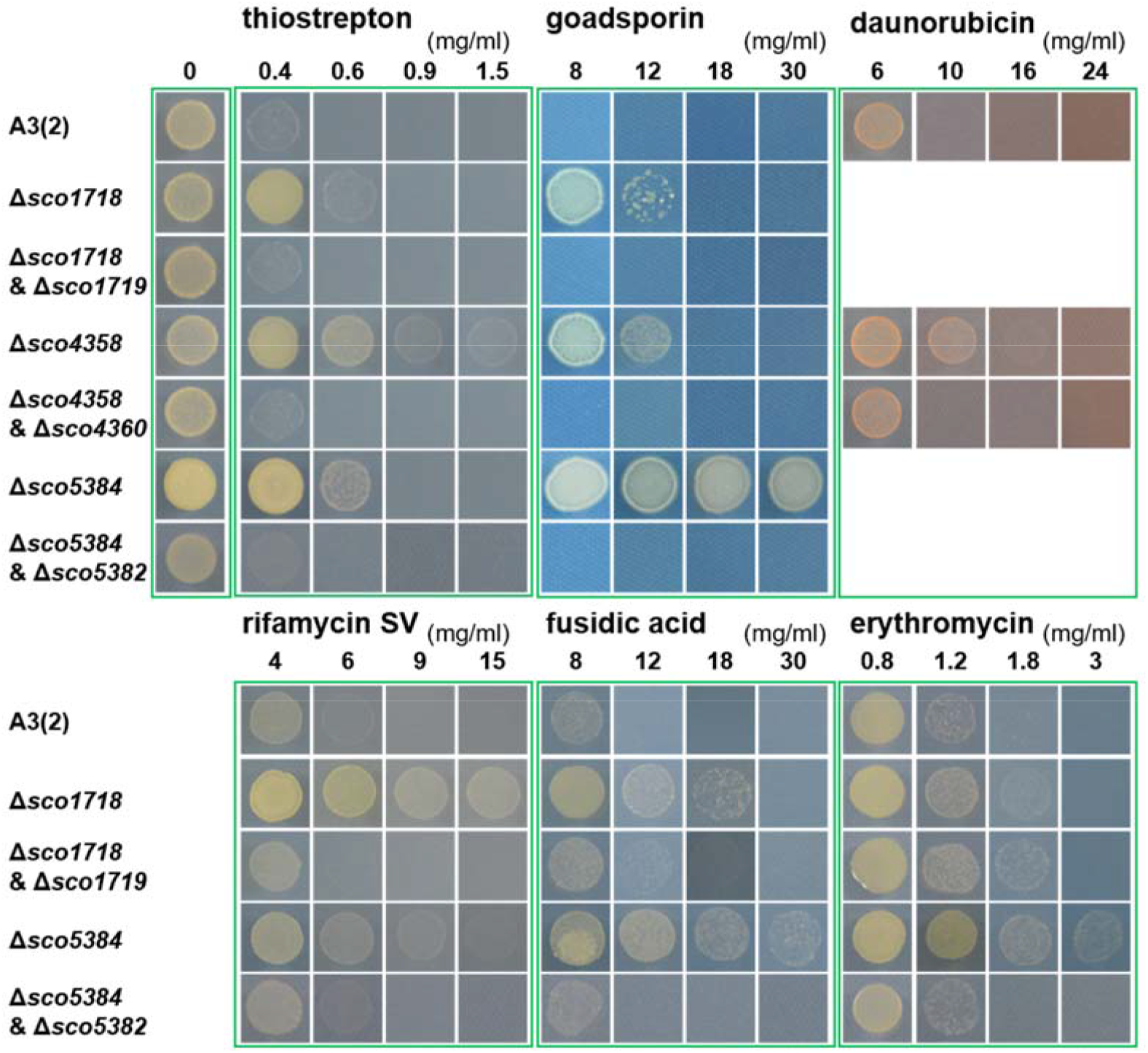
Antibiotic resistance assay on Bennett’s glucose agar. Spore stocks (1 × 10^5^ CFU/ml for each) were spotted on the solid medium containing antibiotics in different concentrations and incubated at 30°C for 2 d.

Additional knockout of ABC transporter genes in the ΔTFR mutants restored the phenotype to that of the A3(2) parent strain, which was sensitive to antibiotics (Fig. 4). These results strongly indicate that the acquired antibiotic resistance of TFR-knockout mutants is the result of ABC transporter overexpression. Notably, based on the range of chemical structures of antibiotics associated with resistance, the three ABC transporters are likely to function as exporters with substrate promiscuity. However, the preference for the substrate was not consistent with molecular size, water solubility (hydrophobic or hydrophilic), and the class of natural products (macrolides and thiopeptides).

### Bioinformatics analysis revealed conservation of TFR-ABC transporter set homologs

To determine the distribution of TFR-ABC transporter sets, we performed a SSDB Gene Cluster Search against the KEGG database (46). Coding sequences showing >100 threshold were chosen (Fig 5, Table S8). The results showed that 114 out of 147 actinobacteria strains were *Streptomyces* spp. and possessed at least one or more homologous sets (65, 92, and 68 out of 147 strains possessed SCO1718-20, SCO4358-60, and SCO5384-82 homologs, respectively). As a142 genome sequences of *Streptomyces* spp. have been deposited in the KEGG database now, result suggest that TFR-ABC transporter sets is relatively distributed (80.3%) in the related species.

**FIG 5.**
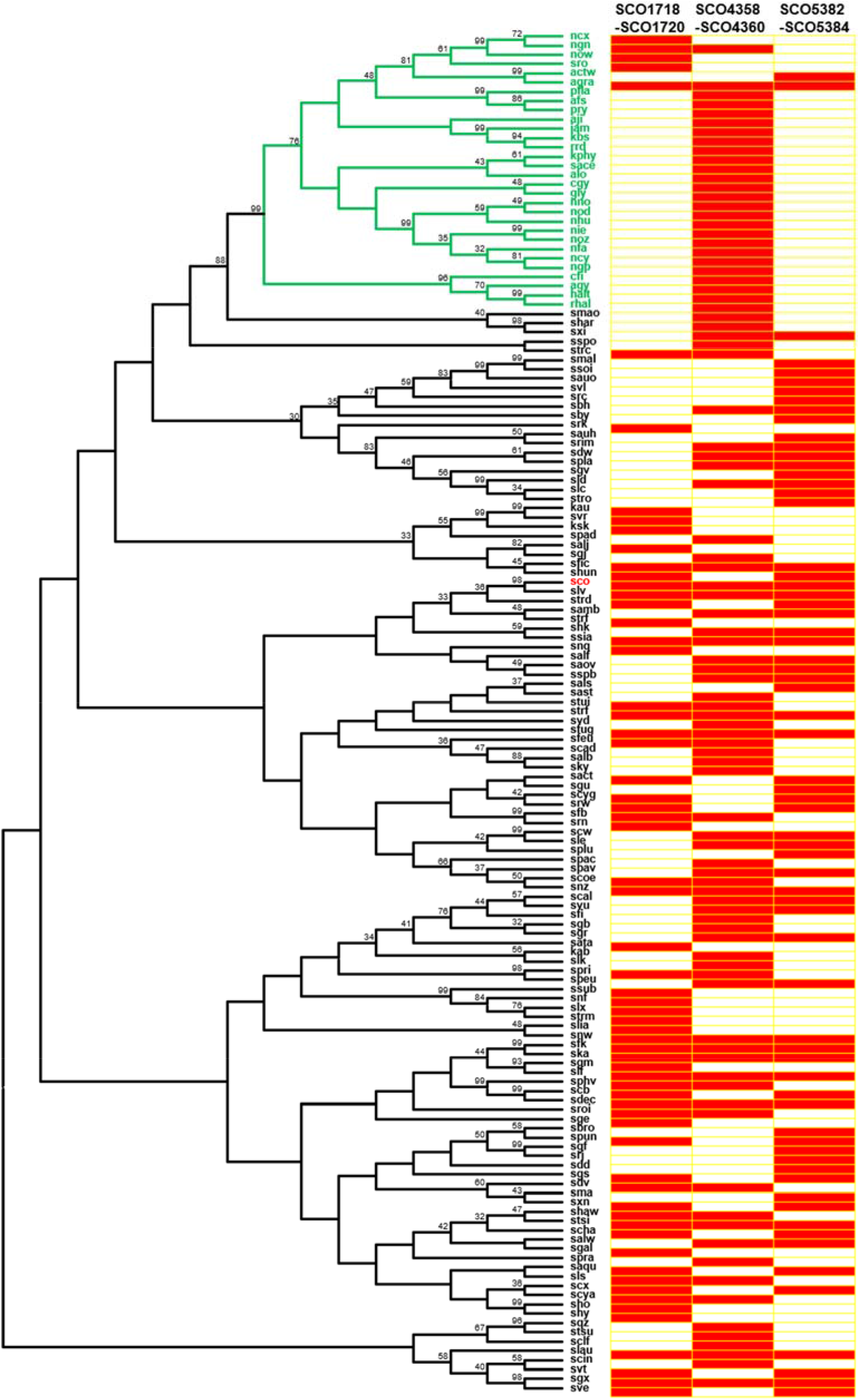
Distribution of TFR-ABC transporter sets in actinomycetes. 16S rDNA sequences were used for maximum likelyhood analysis. Clades in green indicate the non-*Streptomyces* strains. Clades in black indicate the *Streptomyces* strains. Red boxes indicate that conservation of respective TFR-ABC transporter sets.

To evaluate the regulation of physiological expression of ABC transporters, we performed a phylogenetic analysis of SCO1718, SCO4358, and SCO5384 with TFR proteins from *S. coelicolor* A3(2) and other *Streptomyces* species (Tables S9 and S10). SCO1718 and SCO4358 were phylogenetically similar to SimR (47-49), VarR (50), and Pip (SCO4025) (51), and SCO5384 was similar to SlgR1 (52) and SLCG_2919 (53) (Fig. S5). SimR is a well-characterized TFR that represses the expression of the efflux pump SimX in simocyclinone biosynthesis in *Streptomyces antibioticus* Tü 6040 (48). The crystal structure of SimR with its ligand simocyclinone has been solved previously, and the amino acid residues responsible for the ligand binding have been elucidated by Le et al (47).

Multiple sequence alignments were performed for SCO1718, SCO4358, and homologs using ClustalW, and conserved residues were compared with those of SimR (Fig. S6). Notably, although the *N*-terminal regions with the HTH domain responsible for DNA binding were found to be relatively conserved among the tested TFRs, the residues responsible for the binding of simocyclinone in SimR were mostly not conserved among SCO1718, SCO4358, and SimR. These results suggested that SCO1718 and SCO4358 may bind to different ligands for regulation. Further investigation to identify the ligand molecules of TFRs may reveal the physiological functions of the conserved TFR-ABC transporter sets in *Streptomyces* species.

## DISCUSSION

In this study, three sets of TFR-ABC transporters were found to be involved in SM synthesis and multidrug efflux. The expression of ABC transporters has contradicting effects on SM production and sensitivity to natural antibiotics. Previous studies have shown that the expression of the ABC transporters SCO4359-6030 and SCO1719-2029 leads to increased resistance to the antibiotic ADEP (31, 32). In the present study, we further tested the effects of ABC transporter overexpression on SM production and sensitivity to other natural product-origin antibiotics, with another TFR-ABC transporter set, SCO5383-82. The expression of all three ABC transporters regulated by TFRs decreased the production of SMs and increased resistance to natural antibiotics as a common effect (Fig. 6). TFRs are DNA-binding proteins and their binding affinity is generally controlled by specific ligand molecules (34). TFR was originally characterized in the tetracycline efflux system, in which tetracycline binds to TetR and conformational changes induced in TetR result in dissociation from the operator site, allowing the binding of RNA polymerase to the promoter site for the transcription of *tetA* genes encoding MFS-type antiporters, which use H^+^ as a driving force for the efflux of tetracycline (34). It is widely considered that the expression of ABC transporters leads to increased production of SMs and confers self-resistance, since they can reduce the levels of SMs in the cell cytoplasm (33). This includes disruption of *actR-*encoding TFR, leading to overexpression of *actAB*, an RND-type transporter, and results in increased actinorhodin biosynthesis in *S. coelicolor* A3(2) (54). While most studies have focused on the positive effect on the SM production, few studies state that expression of ABC transporters leads to reduced production of SMs; for example, streptomycin production is decreased upon overexpression of the ABC-2 type transporter RspBC in *Streptomyces griseus* (55). However, elucidating the uncharacterized orphan systems that negatively affect SM production is important for a deeper understanding of how the SM production system participates in the whole-cell system. Such cases include genomics studies to select the responsible genes from random mutagenesis in microbial breeding to improve productivity, which has historically been achieved in the fermentation industry. Overexpression of the ABC transporter reduces SM production. One possible explanation for this acquired phenotype is that the physiological substrates that the transporter pumps out are signaling compounds that direct the activation or precursors of SM production in *S. coelicolor* A3(2). *S. coelicolor* A3(2) produces gamma-butyrolactone SCB1 as an elicitor of secondary metabolism for the production of RED, ACT, (56, 57) and cpk (58). Co-culture with wild-type and TFR mutant was performed in the present study; however, we could not detect changes in the apparent phenotype in the wild-type (WT) strain (data not shown). This suggests that transporters have a substrate preference to a certain extent and do not pump out everything, including SCB1. Additionally, overexpression of ABC transporters did not affect the apparent growth.

**FIG 6.**
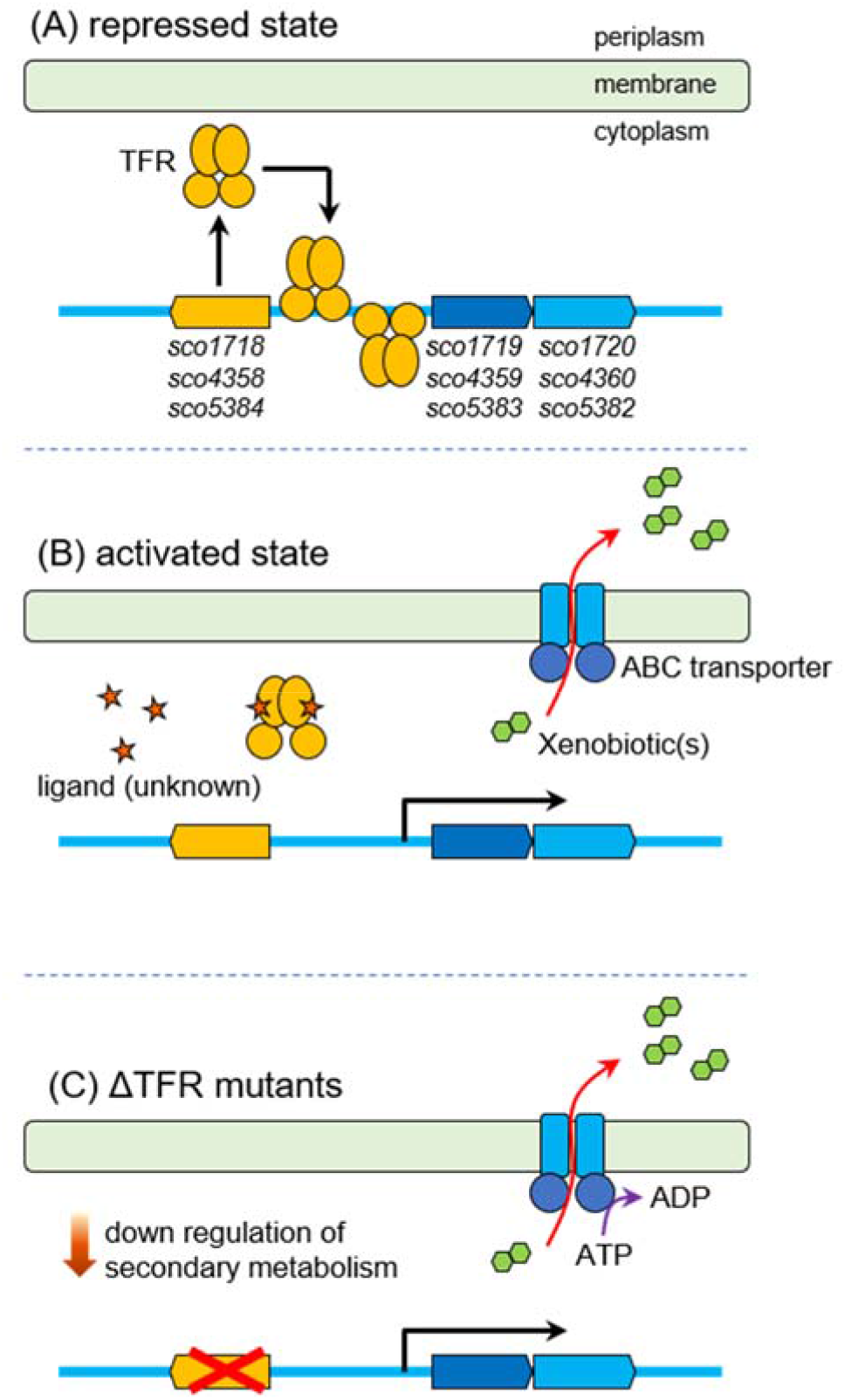
Graphical overview of the TFR-ABC transporter characterized in this study. (A) In the repressed state, each TFRs binds to operator sites and repress the expression of each ABC transporters. (B) Ligand molecule(s) (unknown) bind to the TFRs and facilitate release from the operator sites, which allow expression of each ABC transporter genes. The ABC transporters can efflux xenobiotics from cytoplasm, which confer multidrug resistance. (C) Knockout of TFRs induce overexpression of ABC transporters which consume ATP for xenobiotic efflux, and reduced production of secondary metabolites as well.

It is known that TFR “pseudo” gamma-butyrolactone receptor ScbR2 binds to xenobiotic jadomycin (59, 60). Therefore, we tested whether the 19 antibiotics used in this study could function as ligands for TFRs. However, we did not detect any changes in the shifted bands in the EMSA (data not shown). The TFR-ABC transporter system in this study was standalone, and the ligand molecules for TFRs that regulate the expression of ABC transporters remain obscure. As the expression of ABC transporters (SCO1719-20, SCO4359-60, SCO5383-82) is strictly regulated by dedicated TFRs (SCO1718, SCO4358, SCO5384), further investigation of the ligand molecules that bind to the TFRs will help explain the physiological roles of the conserved ABC transporter system in *Streptomyces*.

TFR two-component ABC transporter sets are widely distributed and conserved among *Streptomyces* species. This implies that acquisition via horizontal gene transfer can spread antibiotic resistance conferred by multidrug efflux pumps. The process of ABC-type ‘active’ drug efflux consumes one molecule of ATP for each conformational change to export substrate. Similarly, the majority of secondary metabolites require ATP for anabolic processes, including polyketide synthase (61) and non-ribosomal peptide synthase (62). Considering the ‘cost’, in other scenarios, these ABC transporters may evolve for the active export of endogenous suppressor molecules of secondary metabolism for other species. Regardless, our results imply the switching of cell metabolism to direct offence (antibiotic production) or defense (efflux pump activation) using costly and limited quantities of cell energy sources (e.g., ATP). These regulatory mechanisms may be advantageous for *Streptomyces* in competitive soil environments.

## MATERIALS AND METHODS

### Bacterial strains and culture conditions

*S. coelicolor* JCM4020 was used in mutagenesis experiments and *S. coelicolor* A3(2) was used for gene knockout and phenotype analysis. *Streptomyces* strains were grown on mannitol soya flour agar medium for sporulation and conjugation and on Bennett’s glucose agar medium for general cultivation. Media used for RED and ACT production in monoculture and combined culture included R2YE medium and YGGS medium [components (in g/l): yeast extract, 3; glucose, 10; glycerol, 20; and soluble starch, 20 (pH 7.2)], respectively (63). NAHU agar medium [components (in g/l): beef extract, 1; yeast extract, 2; peptone, 5; NaCl, 5; histidine, 0.25; uracil, 0.1; and agar, 15 (pH 7.4)], 79NG agar medium [components (in g/l): yeast extract, 2; casamino acids, 2; peptone, 10; NaCl, 6; and agar, 20 (pH 7.3)], and R2YE agar medium were used for CDA, cpk P1, and desferrioxamine production, respectively. *E. coli* strain DH5α was used as the general cloning host, *E. coli* strain ET12567 (pUZ8002) was used as the donor in intergeneric conjugation, and *E. coli* strain BL21 (DE3) was used as the host for heterogeneous protein expression. The *E. coli* strains were grown in Luria broth (LB) supplemented with antibiotics. Apramycin (50 μg/ml), chloramphenicol (25 μg/ml), kanamycin (50 μg/ml), and carbenicillin (100 μg/ml) were added to the growth media as required.

### Gene knockout and gene complementation studies

For *sco1718*, gene knockout was performed via double crossover to generate an in-frame deletion. Briefly, genome fragments corresponding to an in-frame deleted target gene were amplified and ligated into an apramycin resistance-conferring pAT19 plasmid. After introducing the ligated plasmid into the parent strain using the *E. coli* ET12567 (pUZ8002) conjugation method, single crossover exconjugants were selected as apramycin-resistant strains. Subsequently, spores of single crossover exconjugants were re-streaked onto an agar plate, double crossover exconjugants were selected as apramycin-sensitive strains, and the successfully knocked-out strains were verified via colony PCR. For other genes, gene knockout was performed using CRISPR-base editing systems (64) using *S. coelicolor* A3(2) as the parent strain. Briefly, a 20-nucleotide spacer was designed to introduce a nonsense mutation in the gene of interest using CRISPy-web (https://crispy.secondarymetabolites.org/). The plasmid pCRISPR-cBEST, containing a 20-nucleotide spacer, was introduced into the parent strain, and the expression of the base editor based on the Cas9-cytidine deaminase fusion protein was induced by adding thiostrepton. The successfully edited bases were verified using Sanger sequencing. Gene complementation was performed using the pTYM1a integration vector. Briefly, the polycistronic transcription unit containing the gene of interest was amplified using the genomic DNA of the *S. coelicolor* A3(2) strain as a template and ligated into the corresponding site of the plasmid. After introducing the ligated plasmid into the spores of the knockout strain using the *E. coli* ET12567 (pUZ8002) conjugation method, successfully complemented colonies were selected using 25 μg/ml apramycin. *S. coelicolor* A3(2) parent strain and knockout strain that were introduced with a pTYM1a empty vector were also obtained in a similar manner.

### Protein heteroexpression and purification

The transcriptional regulators SCO1718, SCO4358, and SCO5384 were expressed in *E. coli* BL21(DE3) and purified. Briefly, the gene fragments of transcriptional regulators were cloned into the pColdII plasmid (Takara Bio, Kusatsu, Japan) and transformed into *E. coli* BL21(DE3) competent cells. Recombinant *E. coli* BL21(DE3) cells were grown in 100 ml LB medium at 37°C to an optical density at 600 nm of 0.4. Protein expression was induced by adding 0.5 mM isopropyl-β-d-thiogalactoside at 15°C overnight. The proteins were purified using Ni-NTA affinity columns and subsequently subjected to size-exclusion chromatography. The purified protein was confirmed via sodium dodecyl sulphate-polyacrylamide gel electrophoresis and stored at -80°C.

### Electrophoretic mobility shift assay

The palindrome sequences of each intergenic region were obtained as 40-nucleotide oligonucleotides and annealed to obtain DNA probes for EMSA. DNA probes containing mutated palindromic sequences were then obtained. The DNA probes were 3′ end-labelled using digoxigenin (DIG)-11-ddUTP and terminal transferase (Roche, Basel, Switzerland). Binding of TFRs to DNA was analyzed using the DIG Gel Shift Kit, 2nd Generation (Roche). Briefly, 0.1 nM DIG-labelled DNA and varying amounts of transcriptional regulators were mixed and incubated at 16°C for 15 min; the binding reaction mixtures were loaded on 6% (w/v) native polyacrylamide gels and run in TBE buffer at 20 mA for 40 min. After electrophoresis, the gel was electroblotted onto a nylon transfer membrane (GE Healthcare, Chicago, IL, USA) at 300 mA for 60 min and crosslinked with a UV crosslinker at 120 mJ for 3 min. The resulting membrane was incubated in a detection buffer containing anti-digoxigenin-AP Fab fragments. After applying CSPD, chemiluminescence of the membrane was detected using an ImageQuant LAS 4000 system (Cytiva, Marlborough, MA, USA).

### Transcription level analysis via reverse transcription PCR

To analyze the transcription levels of ABC transporter genes, the WT and mutant strains were grown for 24 h in R2YE medium. The cells were harvested from the culture medium and stabilized using the RNAprotect Bacteria Reagent (Qiagen, Hilden, Germany). After mechanical disruption of the cells using the Cell Destroyer PS-1000 (Pro Sense), total RNA was isolated using an RNeasy Mini Kit (Qiagen) following the manufacturer’s protocol. After degradation of DNA using recombinant DNase I (Takara Bio), an equal quantity of total RNA (500 ng) was used in reverse transcription (RT) using PrimeScript RTase (Takara Bio). PCR was performed using GoTaq® Green (Promega, Madison, WI, USA), according to the manufacturer’s protocol for RT. The cDNA fragments corresponding to transcripts of the ABC transporter were detected using SCO1719_RTPCR, SCO4359_RTPCR, and SCO5384_RTPCR primers. The cDNA fragment corresponding to the RNA polymerase principal sigma factor HrdB (SCO5820) transcript was detected using hrdB_RT-PCR primers as a positive control. No signals were detected in the control experiments using RNA samples without RT, indicating that the RT-PCR signals corresponded to the amplification of cDNA transcripts.

### SM assays

For RED and ACT production, *S. coelicolor* A3(2) strains were cultured in R2YE medium at 30°C and culture samples were collected every 24 h. The samples were centrifuged at 12,000 ×g for 5 min, and the absorbance of the supernatant at 640 nm was measured (63). The relative ACT production by each mutant was calculated based on the ratio of the absorbances: A640 mutant/A640 parent strain. The pellets were extracted using methanol for 2 h and the absorbance of the extracted supernatants at 530 nm was measured. The relative RED production in each mutant was calculated using the ratio of absorbances: A530 mutant/A530 parent strain. For CDA production, approximately 1 × 10^5^ spores of each *S. coelicolor* A3(2) strain were spotted on solid NAHU medium and grown at 30°C for 48 h (65). The plates were then overlaid with 12 ml of soft nutrient agar containing 40 mM Ca(NO_3_)_2_ mixed with 120 μl of an overnight culture of *Bacillus subtilis* indicator strain and incubated for 48 h. CDA production was compared based on the size of inhibition zones. A CDA-deficient strain Δ*cdaPS1* was used as the control. For cpk production, approximately 1 × 10^5^ spores of each *S. coelicolor* A3(2) strain were spotted on solid 79NG medium and grown at 30°C for 24 h (66). cpk P1 production was compared based on the appearance of yellow pigments in colonies. For desferrioxamine production, approximately 1 × 10^5^ spores of each *S. coelicolor* A3(2) strain were spotted on solid medium R2YE and grown at 30°C for 48 h (67). The plates were then overlaid with 12 ml of Chrome Azurol S medium (68) without the addition of nutrients. After overnight incubation at 30°C, desferrioxamine production was compared based on the size of orange hollow zones.

### In vivo antibiotic resistance assay

Antibiotic resistance assays were performed using Bennett’s glucose agar supplemented with the indicated concentrations of the compound. Approximately 1 × 10^5^ spores of each *S. coelicolor* A3(2) strain were spotted on each plate, and growth was assessed after incubation at 30°C for 48 h. Antibiotics used to screen for increased resistance are listed in Table S7. The lowest drug concentration that resulted in visual clearing of the agar plates was considered the minimum inhibitory concentration. All experiments were performed in triplicate.

## ACKNOWLEDGMENTS

This research was supported in part by a grant-in-aid from the IFO, Institute for Fermentation, Osaka and the Amano Enzyme Foundation (to H.O. and S.A.); the JSPS A3 Foresight Program (to H.O. and S.A.); JSPS KAKENHI (16K18673 to S.A. and 18H02120 to H.O.); and a general research grant from the IFO, Institute for Fermentation, Osaka (to S.A.). Y. L. was supported by SPRING GX program (https://www.cis-trans.jp/spring_gx/index-e.html). We thank Drs. Katsuya Satoh and Yutaka Oono at the National Institutes for Quantum Science and Technology for carbon-ion beam irradiation for generating the mutants. We thank Masaomi Yanagisawa at The University of Tokyo for the preliminary screening of RED-deficient mutants. We thank Sachiko Kawano at The University of Tokyo for preparing pure goadsporin. We thank Prof. Yasuo Ohnishi and Dr. Takeaki Tezuka at The University of Tokyo for helpful discussions on this research and for providing technical instructions for molecular biology experiments. *S. coelicolor* A3(2) was a kind gift from Prof. Mervin Bibb at John Innes Centre (United Kingdom). The authors declare no conflicts of interest associated with this manuscript.

## CONTRIBUTIONS

YL, SA, and HO designed the study. TI performed the initial genetic experiment on *sco1718*, and YL performed all the other experiments. All authors analyzed and interpreted the data. YL and SA wrote the main manuscript and prepared the figures and tables. HO reviewed the manuscript.

## REFERENCES

1. Barka EA, Vatsa P, Sanchez L, Gaveau-Vaillant N, Jacquard C, Meier-Kolthoff JP, et al. Taxonomy, Physiology, and Natural Products of Actinobacteria (vol 80, pg 1, 2016). Microbiol Mol Biol R. 2016;80(4):Iii–Iii.

2. Newman DJ, Cragg GM. Natural Products as Sources of New Drugs over the Nearly Four Decades from 01/1981 to 09/2019. J Nat Prod. 2020;83(3):770–803.

3. Baltz RH. Gifted microbes for genome mining and natural product discovery. J Ind Microbiol Biot. 2017;44(4-5):573–88.

4. Navarro-Munoz JC, Selem-Mojica N, Mullowney MW, Kautsar SA, Tryon JH, Parkinson EI, et al. A computational framework to explore large-scale biosynthetic diversity. Nat Chem Biol. 2020;16(1):60-+.

5. Rutledge PJ, Challis GL. Discovery of microbial natural products by activation of silent biosynthetic gene clusters. Nat Rev Microbiol. 2015;13(8):509–23.

6. Zarins-Tutt JS, Barberi TT, Gao H, Mearns-Spragg A, Zhang LX, Newman DJ, et al. Prospecting for new bacterial metabolites: a glossary of approaches for inducing, activating and upregulating the biosynthesis of bacterial cryptic or silent natural products. Nat Prod Rep. 2016;33(1):54–72.

7. Van der Meij A, Worsley SF, Hutchings MI, van Wezel GP. Chemical ecology of antibiotic production by actinomycetes. Fems Microbiology Reviews. 2017;41(3):392–416.

8. van der Heul HU, Bilyk BL, McDowall KJ, Seipke RF, van Wezel GP. Regulation of antibiotic production in Actinobacteria: new perspectives from the post-genomic era. Nat Prod Rep. 2018;35(6):575–604.

9. Williamson NR, Fineran PC, Leeper FJ, Salmond GP. The biosynthesis and regulation of bacterial prodiginines. Nat Rev Microbiol. 2006;4(12):887–99.

10. Xu Z, Wang Y, Chater KF, Ou HY, Xu HH, Deng Z, et al. Large-Scale Transposition Mutagenesis of Streptomyces coelicolor Identifies Hundreds of Genes Influencing Antibiotic Biosynthesis. Appl Environ Microbiol. 2017;83(6).

11. Yanagisawa M, Asamizu S, Satoh K, Oono Y, Onaka H. Effects of carbon ion beam-induced mutagenesis for the screening of RED production-deficient mutants of Streptomyces coelicolor JCM4020. Plos One. 2022;17(7):e0270379.

12. Blair JM, Webber MA, Baylay AJ, Ogbolu DO, Piddock LJ. Molecular mechanisms of antibiotic resistance. Nat Rev Microbiol. 2015;13(1):42–51.

13. Du D, Wang-Kan X, Neuberger A, van Veen HW, Pos KM, Piddock LJV, et al. Multidrug efflux pumps: structure, function and regulation. Nat Rev Microbiol. 2018;16(9):523–39.

14. Quistgaard EM, Low C, Guettou F, Nordlund P. Understanding transport by the major facilitator superfamily (MFS): structures pave the way. Nat Rev Mol Cell Biol. 2016;17(2):123–32.

15. Nikaido H. RND transporters in the living world. Res Microbiol. 2018;169(7-8):363–71.

16. Bay DC, Rommens KL, Turner RJ. Small multidrug resistance proteins: a multidrug transporter family that continues to grow. Biochim Biophys Acta. 2008;1778(9):1814–38.

17. Claxton DP, Jagessar KL, McHaourab HS. Principles of Alternating Access in Multidrug and Toxin Extrusion (MATE) Transporters. J Mol Biol. 2021;433(16):166959.

18. Lubelski J, Konings WN, Driessen AJ. Distribution and physiology of ABC-type transporters contributing to multidrug resistance in bacteria. Microbiol Mol Biol Rev. 2007;71(3):463–76.

19. Davidson AL, Dassa E, Orelle C, Chen J. Structure, function, and evolution of bacterial ATP-binding cassette systems. Microbiol Mol Biol Rev. 2008;72(2):317-64, table of contents.

20. Elbourne LD, Tetu SG, Hassan KA, Paulsen IT. TransportDB 2.0: a database for exploring membrane transporters in sequenced genomes from all domains of life. Nucleic Acids Res. 2017;45(D1):D320–D4.

21. Bertram R, Schlicht M, Mahr K, Nothaft H, Saier MH, Jr., Titgemeyer F. In silico and transcriptional analysis of carbohydrate uptake systems of Streptomyces coelicolor A3(2). J Bacteriol. 2004;186(5):1362–73.

22. Martin JF, Liras P. Molecular Mechanisms of Phosphate Sensing, Transport and Signalling in Streptomyces and Related Actinobacteria. Int J Mol Sci. 2021;22(3).

23. Iinuma C, Saito A, Ohnuma T, Tenconi E, Rosu A, Colson S, et al. NgcE(Sco) Acts as a Lower-Affinity Binding Protein of an ABC Transporter for the Uptake of N,N’-Diacetylchitobiose in Streptomyces coelicolor A3(2). Microbes Environ. 2018;33(3):272–81.

24. Saito A, Shinya T, Miyamoto K, Yokoyama T, Kaku H, Minami E, et al. The dasABC gene cluster, adjacent to dasR, encodes a novel ABC transporter for the uptake of N,N’-diacetylchitobiose in Streptomyces coelicolor A3(2). Appl Environ Microbiol. 2007;73(9):3000–8.

25. Barona-Gomez F, Lautru S, Francou FX, Leblond P, Pernodet JL, Challis GL. Multiple biosynthetic and uptake systems mediate siderophore-dependent iron acquisition in Streptomyces coelicolor A3(2) and Streptomyces ambofaciens ATCC 23877. Microbiology (Reading). 2006;152(Pt 11):3355–66.

26. Lautru S, Deeth RJ, Bailey LM, Challis GL. Discovery of a new peptide natural product by Streptomyces coelicolor genome mining. Nat Chem Biol. 2005;1(5):265–9.

27. Hojati Z, Milne C, Harvey B, Gordon L, Borg M, Flett F, et al. Structure, biosynthetic origin, and engineered biosynthesis of calcium-dependent antibiotics from Streptomyces coelicolor. Chem Biol. 2002;9(11):1175–87.

28. Kodani S, Hudson ME, Durrant MC, Buttner MJ, Nodwell JR, Willey JM. The SapB morphogen is a lantibiotic-like peptide derived from the product of the developmental gene ramS in Streptomyces coelicolor. Proc Natl Acad Sci U S A. 2004;101(31):11448–53.

29. O’Connor TJ, Nodwell JR. Pivotal roles for the receiver domain in the mechanism of action of the response regulator RamR of Streptomyces coelicolor. J Mol Biol. 2005;351(5):1030–47.

30. Hesketh A, Kock H, Mootien S, Bibb M. The role of absC, a novel regulatory gene for secondary metabolism, in zinc-dependent antibiotic production in Streptomyces coelicolor A3(2). Mol Microbiol. 2009;74(6):1427–44.

31. Compton CL, Carney DW, Groomes PV, Sello JK. Fragment-Based Strategy for Investigating and Suppressing the Efflux of Bioactive Small Molecules. Acs Infect Dis. 2015;1(1):53–8.

32. Gominet M, Seghezzi N, Mazodier P. Acyl depsipeptide (ADEP) resistance in Streptomyces. Microbiology (Reading). 2011;157(Pt 8):2226–34.

33. Cuthbertson L, Nodwell JR. The TetR Family of Regulators. Microbiol Mol Biol R. 2013;77(3):440–75.

34. Ramos JL, Martinez-Bueno M, Molina-Henares AJ, Teran W, Watanabe K, Zhang X, et al. The TetR family of transcriptional repressors. Microbiol Mol Biol Rev. 2005;69(2):326–56.

35. Getsin I, Nalbandian GH, Yee DC, Vastermark A, Paparoditis PC, Reddy VS, et al. Comparative genomics of transport proteins in developmental bacteria: Myxococcus xanthus and Streptomyces coelicolor. BMC Microbiol. 2013;13:279.

36. Asamizu S, Ozaki T, Teramoto K, Satoh K, Onaka H. Killing of Mycolic Acid-Containing Bacteria Aborted Induction of Antibiotic Production by Streptomyces in Combined-Culture. Plos One. 2015;10(11).

37. Kato M, Asamizu S, Onaka H. Intimate relationships among actinomycetes and mycolic acid-containing bacteria. Sci Rep-Uk. 2022;12(1).

38. Onaka H, Mori Y, Igarashi Y, Furumai T. Mycolic Acid-Containing Bacteria Induce Natural-Product Biosynthesis in Streptomyces Species. Appl Environ Microb. 2011;77(2):400–6.

39. Asamizu S, Pramana AAC, Kawai SJ, Arakawa Y, Onaka H. Comparative Metabolomics Reveals a Bifunctional Antibacterial Conjugate from Combined-Culture of Streptomyces hygroscopicus HOK021 and Tsukamurella pulmonis TP-B0596. Acs Chem Biol. 2022;17(9):2664–72.

40. Hoshino S, Onaka H, Abe I. Activation of silent biosynthetic pathways and discovery of novel secondary metabolites in actinomycetes by co-culture with mycolic acid-containing bacteria. J Ind Microbiol Biotechnol. 2019;46(3-4):363–74.

41. Gilchrist CLM, Chooi YH. Clinker & clustermap.js: Automatic generation of gene cluster comparison figures. Bioinformatics. 2021.

42. Kumar S, Stecher G, Li M, Knyaz C, Tamura K. MEGA X: Molecular Evolutionary Genetics Analysis across Computing Platforms. Mol Biol Evol. 2018;35(6):1547–9.

43. Gomez-Escribano JP, Song LJ, Fox DJ, Yeo V, Bibb MJ, Challis GL. Structure and biosynthesis of the unusual polyketide alkaloid coelimycin P1, a metabolic product of the cpk gene cluster of Streptomyces coelicolor M145. Chem Sci. 2012;3(9):2716–20.

44. Barona-Gomez F, Wong U, Giannakopulos AE, Derrick PJ, Challis GL. Identification of a cluster of genes that directs desferrioxamine biosynthesis in Streptomyces coelicolor M145. J Am Chem Soc. 2004;126(50):16282–3.

45. Onaka H, Nakaho M, Hayashi K, Igarashi Y, Furumai T. Cloning and characterization of the goadsporin biosynthetic gene cluster from Streptomyces sp. TP-A0584. Microbiology (Reading). 2005;151(Pt 12):3923–33.

46. Kanehisa M, Furumichi M, Tanabe M, Sato Y, Morishima K. KEGG: new perspectives on genomes, pathways, diseases and drugs. Nucleic Acids Res. 2017;45(D1):D353–D61.

47. Le TB, Stevenson CE, Fiedler HP, Maxwell A, Lawson DM, Buttner MJ. Structures of the TetR-like simocyclinone efflux pump repressor, SimR, and the mechanism of ligand-mediated derepression. J Mol Biol. 2011;408(1):40–56.

48. Le TB, Fiedler HP, den Hengst CD, Ahn SK, Maxwell A, Buttner MJ. Coupling of the biosynthesis and export of the DNA gyrase inhibitor simocyclinone in Streptomyces antibioticus. Mol Microbiol. 2009;72(6):1462–74.

49. Le TB, Schumacher MA, Lawson DM, Brennan RG, Buttner MJ. The crystal structure of the TetR family transcriptional repressor SimR bound to DNA and the role of a flexible N-terminal extension in minor groove binding. Nucleic Acids Res. 2011;39(21):9433–47.

50. Namwat W, Lee CK, Kinoshita H, Yamada Y, Nihira T. Identification of the varR gene as a transcriptional regulator of virginiamycin S resistance in Streptomyces virginiae. J Bacteriol. 2001;183(6):2025–31.

51. Folcher M, Morris RP, Dale G, Salah-Bey-Hocini K, Viollier PH, Thompson CJ. A transcriptional regulator of a pristinamycin resistance gene in Streptomyces coelicolor. J Biol Chem. 2001;276(2):1479–85.

52. Gomez C, Olano C, Mendez C, Salas JA. Three pathway-specific regulators control streptolydigin biosynthesis in Streptomyces lydicus. Microbiology (Reading). 2012;158(Pt 10):2504–14.

53. Xu Y, Ke M, Li J, Tang Y, Wang N, Tan G, et al. TetR-Type Regulator SLCG_2919 Is a Negative Regulator of Lincomycin Biosynthesis in Streptomyces lincolnensis. Appl Environ Microbiol. 2019;85(1).

54. Xu Y, Willems A, Au-Yeung C, Tahlan K, Nodwell JR. A two-step mechanism for the activation of actinorhodin export and resistance in Streptomyces coelicolor. mBio. 2012;3(5):e00191–12.

55. Takano H, Toriumi N, Hirata M, Amano T, Ohya T, Shimada R, et al. An ABC transporter involved in the control of streptomycin production in Streptomyces griseus. FEMS Microbiol Lett. 2016;363(14).

56. Takano E, Nihira T, Hara Y, Jones JJ, Gershater CJ, Yamada Y, et al. Purification and structural determination of SCB1, a gamma-butyrolactone that elicits antibiotic production in Streptomyces coelicolor A3(2). J Biol Chem. 2000;275(15):11010–6.

57. Takano E, Chakraburtty R, Nihira T, Yamada Y, Bibb MJ. A complex role for the gamma-butyrolactone SCB1 in regulating antibiotic production in Streptomyces coelicolor A3(2). Mol Microbiol. 2001;41(5):1015–28.

58. Takano E, Kinoshita H, Mersinias V, Bucca G, Hotchkiss G, Nihira T, et al. A bacterial hormone (the SCB1) directly controls the expression of a pathway-specific regulatory gene in the cryptic type I polyketide biosynthetic gene cluster of Streptomyces coelicolor. Mol Microbiol. 2005;56(2):465–79.

59. Wang W, Ji J, Li X, Wang J, Li S, Pan G, et al. Angucyclines as signals modulate the behaviors of Streptomyces coelicolor. Proc Natl Acad Sci U S A. 2014;111(15):5688–93.

60. Xu G, Wang J, Wang L, Tian X, Yang H, Fan K, et al. “Pseudo” gamma-butyrolactone receptors respond to antibiotic signals to coordinate antibiotic biosynthesis. J Biol Chem. 2010;285(35):27440–8.

61. Khosla C, Herschlag D, Cane DE, Walsh CT. Assembly line polyketide synthases: mechanistic insights and unsolved problems. Biochemistry-Us. 2014;53(18):2875–83.

62. Walsh CT. Insights into the chemical logic and enzymatic machinery of NRPS assembly lines. Nat Prod Rep. 2016;33(2):127–35.

63. Kieser T, Bibb MJ, Buttner MJ, Chater KF, Hopwood DA. Practical Streptomyces Genetics: John Innes Foundation; 2000.

64. Tong Y, Whitford CM, Robertsen HL, Blin K, Jorgensen TS, Klitgaard AK, et al.Highly efficient DSB-free base editing for streptomycetes with CRISPR-BEST. Proc Natl Acad Sci U S A. 2019;116(41):20366–75.

65. Chong PP, Podmore SM, Kieser HM, Redenbach M, Turgay K, Marahiel M, et al. Physical identification of a chromosomal locus encoding biosynthetic genes for the lipopeptide calcium-dependent antibiotic (CDA) of Streptomyces coelicolor A3(2). Microbiology (Reading). 1998;144 (Pt 1):193–9.

66. Pawlik K, Kotowska M, Kolesinski P. Streptomyces coelicolor A3(2) produces a new yellow pigment associated with the polyketide synthase Cpk. J Mol Microbiol Biotechnol. 2010;19(3):147–51.

67. Craig M, Lambert S, Jourdan S, Tenconi E, Colson S, Maciejewska M, et al. Unsuspected control of siderophore production by N-acetylglucosamine in streptomycetes. Environ Microbiol Rep. 2012;4(5):512–21.

68. Schwyn B, Neilands JB. Universal chemical assay for the detection and determination of siderophores. Anal Biochem. 1987;160(1):47–56.

